# Seeing the World Like Never Before: Human stereovision through perfect optics

**DOI:** 10.1101/2021.01.05.425427

**Authors:** Cherlyn J Ng, Randolph Blake, Martin S Banks, Duje Tadin, Geunyoung Yoon

## Abstract

Stereovision is the ability to perceive fine depth variations from small differences in the two eyes’ images. Using adaptive optics, we show that even minute optical aberrations that are not clinically correctable, and go unnoticed in everyday vision, can affect stereo acuity. Hence, the human binocular system is capable of using unnaturally fine details that are not encountered in everyday vision. More importantly, stereoacuity was still considerably variable even with perfect optics. This variability can be attributed to neural adaptation. Our visual system tries to compensate for these aberrations through neural adaptation that optimizes stereovision when viewing stimuli through one’s habitual optics. However, the same adaptation becomes ineffective when the optics are changed, even if improved. Beyond optical imperfections, we show that stereovision is limited by neural adaptation to one’s own optics.

**Significance statement:** Humans, and animals with front-facing eyes, view the world from slightly different vantage points. This creates small differences in the left and right images that can be utilized for fine depth perception (stereovision). Retinal images are also subject to imperfections that are often different in the optics of the two eyes. Using advanced optical correction techniques, we show that even the smallest imperfections that escape clinical detection affect stereovision. We also find that neural processes become adapted to a person’s own optics. Hence, stereovision is directly impacted by the optics of the eyes, and indirectly via neural adaptation. Since the optics change over the lifespan, our results imply that the adult binocular system is adaptable with possibilities for binocular rehabilitation.

## Introduction

Many of us have refractive errors that degrade the quality of the images formed on our retinas, the most common errors being myopia, hyperopia, and astigmatism. If uncorrected, these optical defects can compromise everyday activities such as visually guided behavior and reading. Collectively known as lower-order aberrations, these defects are easily corrected with spectacles or contact lenses. In addition, we all have other optical aberrations that cannot be so easily corrected. These defects---higher-order aberrations---also degrade retinal images [1]. We are not aware of these residual native aberrations because our everyday experience provides no basis for knowing what the world would look like if those aberrations were eliminated. This conundrum raises a fascinating, albeit modest version of Molyneux’s problem [2]: What would the world look like if a person were able to see the world through eyes with perfect optics? Answering this question will specify the degree to which the higher-order aberrations, as well as the small amounts of residual lower-order ones, limit human visual function and, in turn, reveal the extent to which the visual nervous system can utilize spatial information never before encountered.

This question can be answered using adaptive optics (AO). AO is a technique that measures optical wavefront distortions through the pupil of the eye and compensates by setting a complementary shape on a deformable mirror that reflects visual stimuli into the eye to achieve near perfect, diffraction-limited retinal images [3]. Previous work has shown that removing higher-order aberrations yields significant improvements in visual acuity and contrast sensitivity [4-7]. This previous work dealt with monocular vision but humans are intrinsically binocular and the question remains as to how aberrations of both eyes affect binocular vision. We utilized AO to examine human stereopsis, an aspect of binocular vision that exploits tiny positional differences in the two eyes’ retinal images [8-10]. These positional differences---binocular disparities---are estimated in visual cortex and produce a compelling sensation of three dimensionality [11, 12]. As precise stereopsis derives from small differences between the two eyes, it may be especially susceptible to imperfections associated with optical aberrations, especially higher-order ones, because those aberrations typically differ in the two eyes.

Stereopsis, like other visual functions, is adversely affected by blur [13-15], particularly when the images to the two eyes differ in optical quality [16-18]. It is not uncommon that the two eyes have different spherical refractive errors (anisometropia and monovision are examples [19, 20]) and astigmatism of different magnitudes and axis. Even in well-focused eyes, higher-order aberration profiles are seldom the same in the two eyes [21].

Are there consequences of living with chronic and conventionally uncorrectable aberrations? We know that monocular images appear sharpest when they are presented with a person’s native aberrations rather than other aberrations of the same magnitude. This observation strongly suggests that people adapt to the blur caused by their own optics [22-24]. Binocular adaptation to habitual optics also biases the cyclopean percept of blurriness [25]. Plausibly, stereopsis would capitalize on such adaptation too and thereby sharpen depth perception under habitually experienced conditions. This explanation was cited in [26] as a reason for not observing improvement in stereo vision when optical aberrations were corrected.

With these points in mind, we investigated whether stereopsis is limited by the optics of the two eyes, and, as a follow up, whether the neural mechanisms underlying stereopsis become adapted to an individual’s degraded retinal images. We pursued this by measuring stereoacuity when all eye aberrations were eliminated by AO correction, in comparison to measurements with native optics when the familiar lower and higher-order aberrations were in place. Improvement in AO-assisted stereopsis would be noteworthy because it would show that the brain can exploit greater image sharpness than ever experienced before (Figure 1).

**Figure 1.**
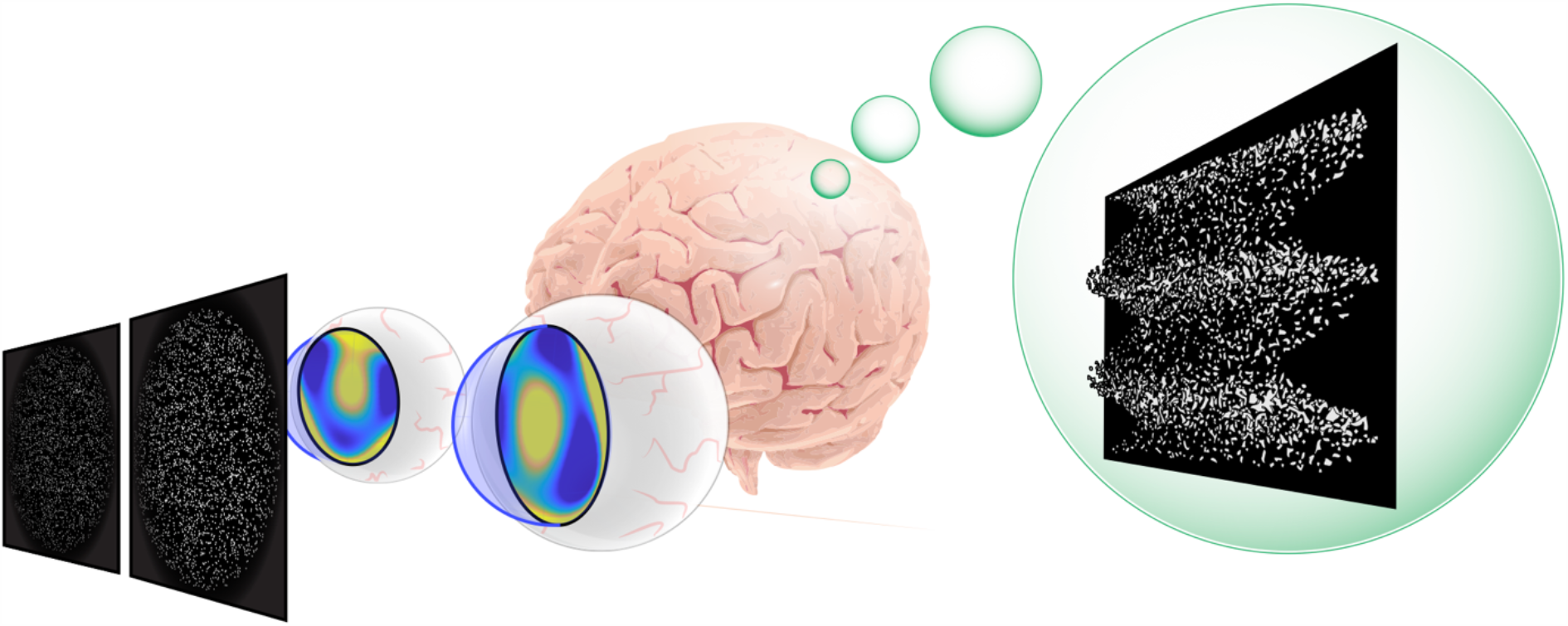
Schematic of the paradigm. The aim of the study was to elucidate the interplay between optics and neural adaptation in determining stereoacuity.

## Results

### Visual acuity follows optical quality

We first wanted to confirm that correcting the optical aberrations using AO yields improvement in monocular visual acuity where improvement was expected. We focused on monocular letter acuity because others had shown significant improvement in that task when aberrations are corrected by AO [4-6, 27].

Figure 2A shows examples of an individual’s aberrated wavefront pattern, and the AO-corrected pattern and the associated retinal images. Figure 2B plots visual acuity with and without AO correction. Note that native optics in our study always included the best conventional refractive correction (sphere and cylinder), if needed. The average acuities for the left eye improved with AO correction from 20/18 to 20/12.3 (an improvement of 31.7%) and for the right eye from 20/17 to 20/11 (35.3% improvement). The improvements were statistically significant (t_13_ = 10.0, p < 10^−7^, one tailed, all eyes combined). Indeed, the corrected acuities approached limits set by photoreceptor spacing at the fovea [28, 29]. Participants reported that stimuli viewed with AO correction appeared unusually and surprisingly sharp. These results confirm that our AO correction substantially improves retinal-image quality.

**Figure 2:**
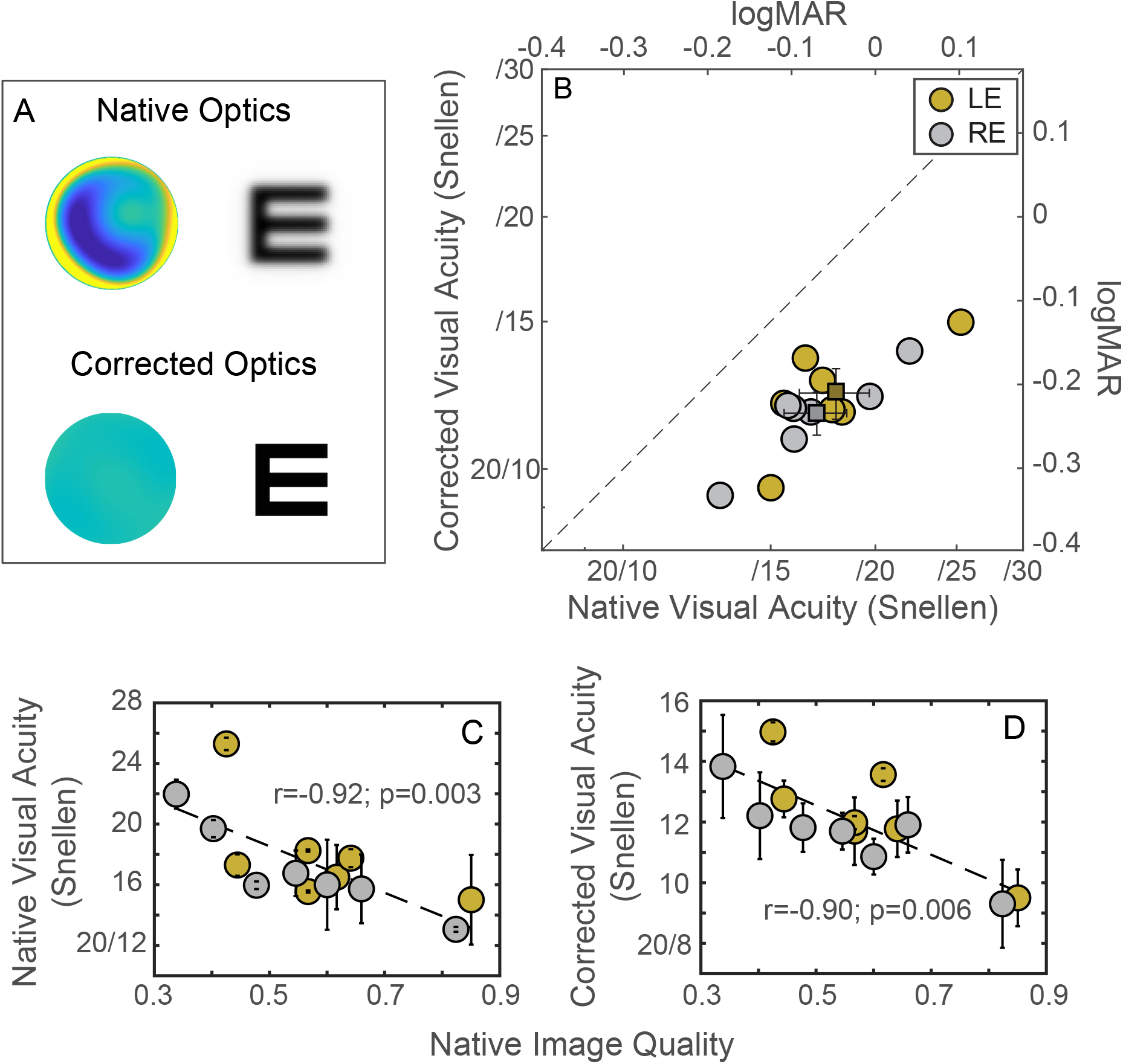
Visual acuity and optical correction. (A) Wavefronts and retinal images from the right eye of a representative participant (S6). The left panels are maps of the wavefront drawn in the same color scale in the native and corrected conditions. The color variation represents the distortion of the wavefront. The right panels are simulations of the retinal images of a 20/20 Snellen letter E in the two conditions. (B) Monocular visual acuity with native and corrected optics. Visual acuity with AO-corrected optics is plotted against acuity with native optics for both eyes of all participants. Gold and silver symbols indicate the acuities for the left and right eyes, respectively. The left and bottom axes are acuity in Snellen notation. The right and top axes are the equivalent in logMAR units (logarithm of minimal angle of resolution in minutes of arc). The small squares are the average values for the left and right eyes. (C) Native and (D) corrected visual acuity as a function of native retinal-image quality, simulated by convolving individual eyes’ PSFs with a 20/20 Snellen E (see Methods). All error bars indicate ±1SD.

Visual acuity with the native optics was highly correlated with image quality (Figure 2C; Pearson r_12_ = −0.92; p = 0.003). This was entirely expected as it simply shows that those individuals with better native optics have better visual acuity. Surprisingly, we also found a similar degree of individual variability in the acuities measured under AO correction even though participants had essentially the same (near-perfect) image quality in that condition (Figures 2B and S1). Notably, the acuities with corrected optics were correlated with image quality before AO correction (Figure 2D; Pearson r_12_ = −0.90; p = 0.006): specifically, those with poor native optics also exhibited relatively poor visual acuity under AO correction. We examine the implications of these intriguing observations in the Discussion.

### Stereo acuity improves with optical correction

We next turned to the main topic of investigation: How do the eyes’ optics affect the precision of stereopsis? To answer that question, we measured the smallest disparity that allowed participants to identify the orientation of a disparity-defined depth corrugation ([30, 31]; Equations 1 and 2 in the Methods) with their native optics and with AO-corrected optics. Stereo acuity was measured at three corrugation frequencies: 1, 2, and 3cpd.

As can be seen in Figure 3B, there was a clear improvement in stereoacuity with AO correction at all three corrugation frequencies. The average improvement (small squares with error bars) was 30.0%, a statistically significant improvement (F_1,36_=4.08, p=0.050; two-way repeated randomized-block ANOVA) and similar in magnitude to the improvement in visual acuity with AO correction (Figure 2). Stereo acuity increased with increasing corrugation frequency (F_2,36_=14.96, p<10^−4^), but the improvement with AO-corrected optics relative to native optics was similar at all frequencies (frequency x correction interaction: F_2,36_=0.21, p=0.81).

**Figure 3:**
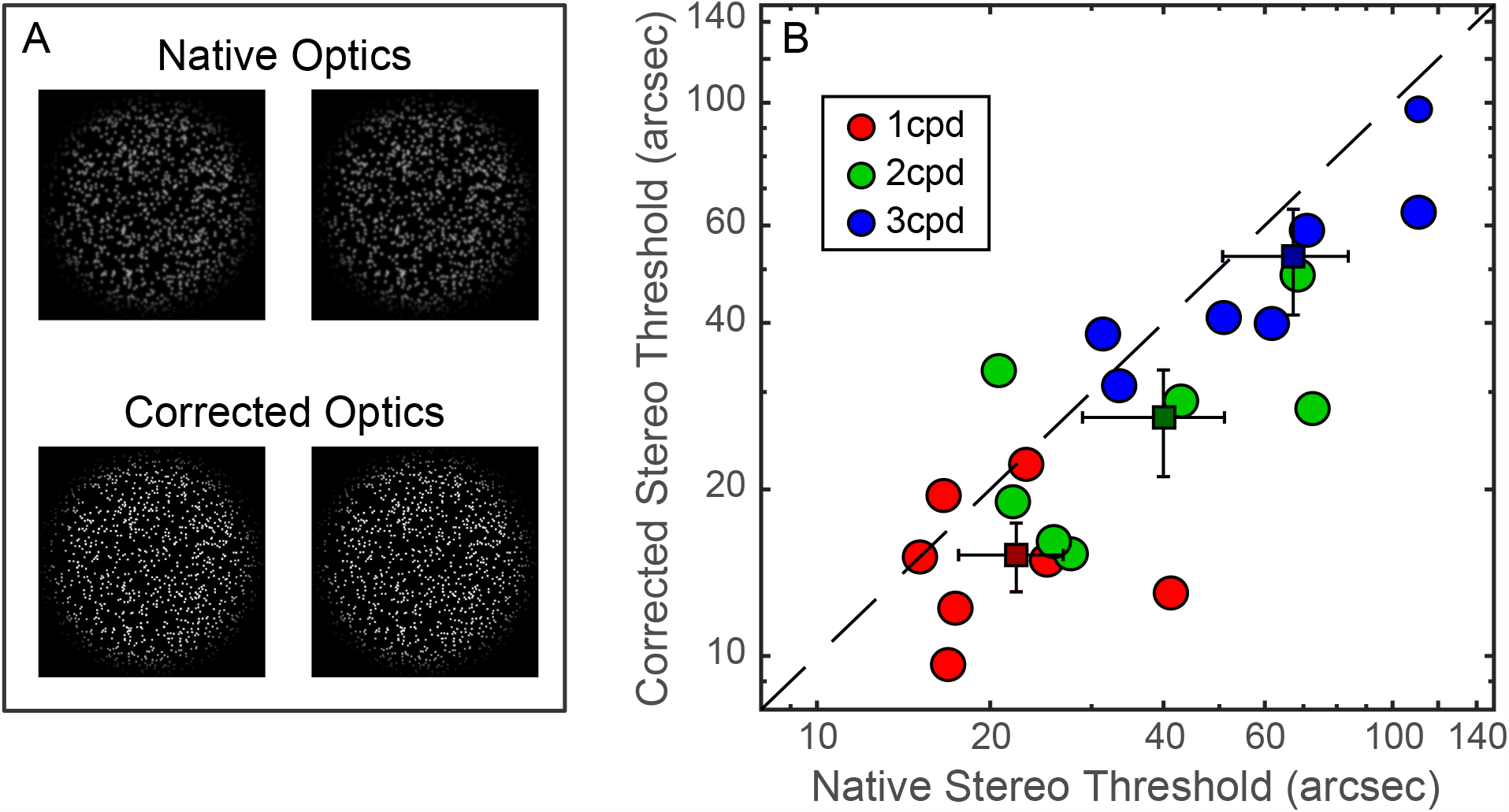
Stereo thresholds and optical correction. **(A)** Simulated retinal images, one for each eye of the random-dot stimulus, are depicted for the native- and AO-corrected-optics conditions. The reader can cross fuse to observe the depth corrugation. (B) Stereo thresholds with native and AO-corrected optics. The smallest discriminable disparity with correction is plotted against the smallest discriminable disparity with native optics for each participant and corrugation frequency (red for 1cpd, green for 2cpd, and blue for 3cpd). The small squares are the average values for each frequency. Errors bars are standard deviations.

Most participants did substantially better in the stereo test with corrected optics. However, as with visual acuity (Figure 2D), we found considerable individual differences in AO-corrected stereo acuity even though retinal-image quality was essentially the same for all of them in that condition. This includes one participant who performed slightly worse with AO correction than with native optics (the three data points above the identity line), which we will address later.

### What causes individual differences in stereo threshold?

Previous studies have found significant individual differences in stereo acuity [32, 33]; we observed this with native optics (horizontal spread in Figure 3B), too. Why do people differ in this task? To tackle that question, we investigated whether peoples’ native optical aberrations determined the individual differences. There are two possibilities. 1) Some people simply have better optics than others. Consequently, those with minimal higher-order aberrations, and hence better image quality, are able to perform better. This hypothesis is consistent with the observation that stereo acuity is better with sharp images in the two eyes than with blurred images [14, 34]. Our results show this too because stereo acuity with AO correction was generally better than acuity without AO correction (Figure 3B). 2) Alternatively, individual differences in stereo acuity might largely derive from interocular differences in the aberrations. This hypothesis is suggested by the *blur paradox*: i.e., stereo acuity is actually better when both eyes’ images are equivalently blurred compared to when only one eye’s image is blurred and the other eye’s image is not blurred. The blur paradox implies that the binocular matching required to see depth from disparity is dependent on having images of equivalent contrast energy and spatial-frequency content in the two eyes. We next sought to determine which of the two hypotheses is the better predictor of stereo acuity across individuals.

We simulated what the left and right retinal images for each participant would be by convolving their PSFs with our random-dot stimuli. From the resulting images, we quantified binocular image quality in two ways: the average image quality (*ImQ*_*Mean*_– Equation 4) and the inter-ocular difference in image quality (*ImQ*_*IOD*;_ - Equation 5; see Methods). Results from those simulations are summarized in Figure 4.

**Figure 4:**
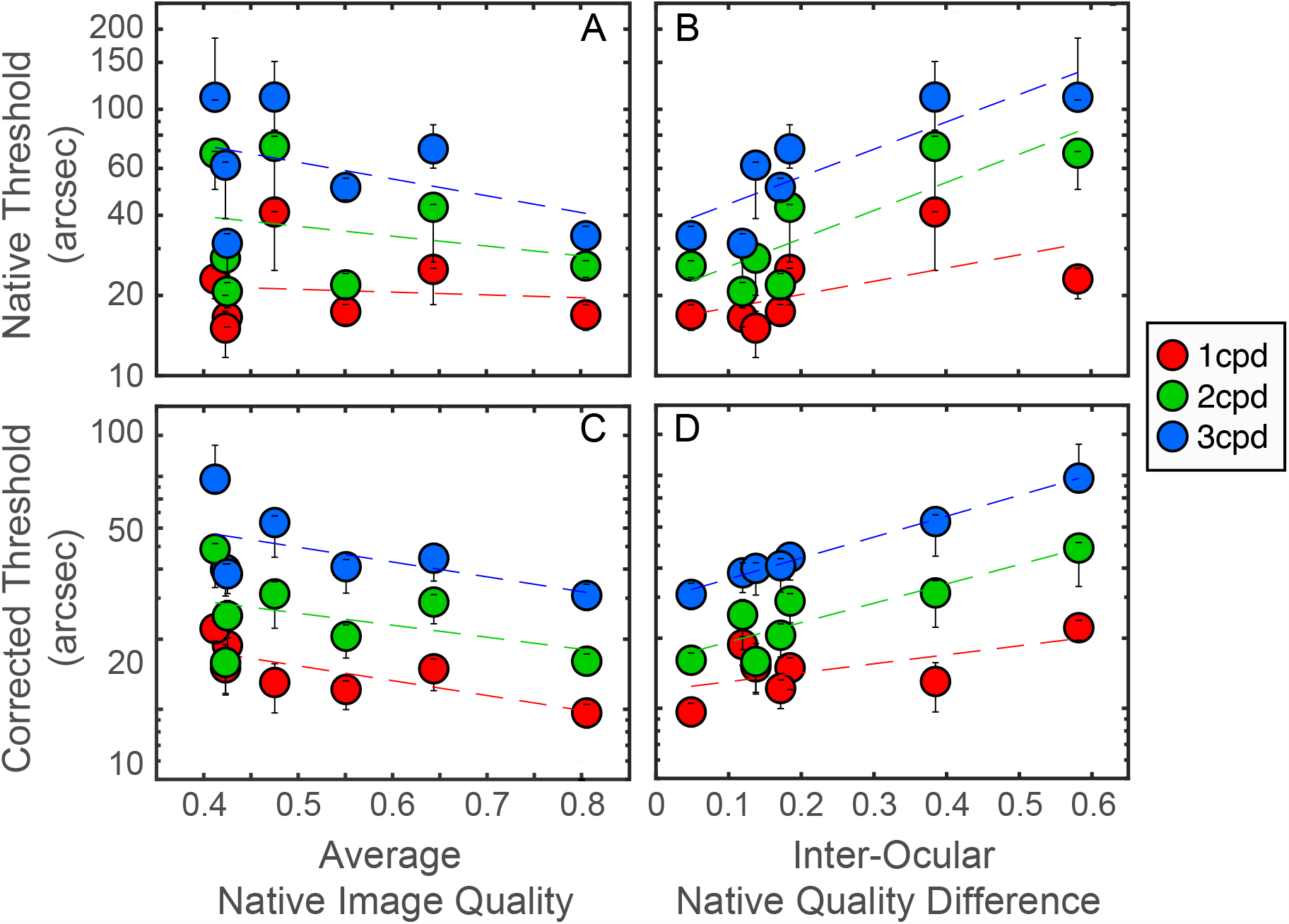
Stereo thresholds and image quality. (A and B) Stereo threshold with native optics as a function of the average (*ImQMean* – Equation 4; A) and inter-ocular difference (*ImQIOD* – Equation 5; B) in the native image quality. The smallest discriminable disparity was plotted for each participant and corrugation frequency against the average image quality for the two eyes. Datapoints for each participant were aligned vertically. Error bars indicate ±1SD. (C and D) Stereo threshold with AO-corrected optics as a function of the same.

Figure 4A plots stereo acuity with native optics as a function of average image quality; they do not covary systematically (Pearson’s r_5_= −0.13, −0.30, −0.43 at 1, 2, and 3cpd; all p > 0.34). Figure 4B plots native stereo acuity as a function of inter-ocular difference in quality. Here we see a clear association: There were significant positive correlations for corrugation frequencies of 2 and 3cpd (Pearson r_5_ = 0.89 and 0.91 respectively, p<0.01 in both cases) and a positive trend for 1cpd (Pearson r_5_ = 0.55, p=0.13). In other words, participants with approximately equal native optical quality in the two eyes exhibit better stereo acuity than participants with larger interocular differences. These results reveal that the blur paradox is not specific to major defocus, but generalizes to smaller interocular differences caused higher-order aberrations.

Individual differences in stereo acuity under native optics were expected. But we also found similarly large inter-subject variability under AO correction (vertical spread in Figure 3B). That is, there were still notable individual differences when optical quality was essentially the same in all eyes because the optical imperfections had been corrected. Interestingly, participants’ stereo acuity with AO-corrected optics was significantly correlated with their stereo acuity with their native optics at the higher corrugation frequencies (1cpd: Pearson’s r_5_ = −0.048, p= 0.92; 2cpd: r_5_= 0.80, p= 0.032; 3cpd: r_5_= 0.86, p= 0.013).

Similar to our observations of stereo acuity with native optics, overall image quality was not significantly correlated with AO performance. (Figure 4C; 1cpd: Pearson’s r_5_ = − 0.73, p= 0.06; 2cpd: r_5_= −0.43, p= 0.33; 3cpd: r_5_ = −0.50; p= 0.25). But inter-ocular difference in native image quality was well correlated with AO-corrected stereo acuity even though the optical aberrations had been eliminated (Figure 4D; 1cpd: Pearson’s r_5_ = 0.63, p= 0.13; 2cpd: r_5_ = 0.93, p= 0.0024; 3cpd: r_5_= 0.99, p< 0.0001).

In summary, individuals who have had large interocular differences in optical quality during everyday viewing had poorer stereo acuity with their native optics but also with AO-corrected optics (i.e., when the image quality in the two eyes was essentially the same for all participants; Figure S1). The finding with corrected optics is paradoxical because the people who received the greatest improvement in inter-ocular difference, had the worst stereo acuity with AO-corrected optics. Why would people with larger inter-ocular differences have poorer stereopsis than people with smaller differences when the differences have been eliminated? We explore that question next.

### Adaptation to native optics

Our results show that individuals with poorer native optics (specifically, larger differences between the eyes) exhibit poorer stereo acuity even when their optics are corrected and equated in the two eyes. This inability to achieve greater improvement in stereo acuity implies that neural circuits subserving stereopsis have been shaped by the visual experience delimited by their native optics. To test this hypothesis further, we replaced the optics of one person with those of another and measured the effect on stereo acuity (Figure 5A).

**Figure 5:**
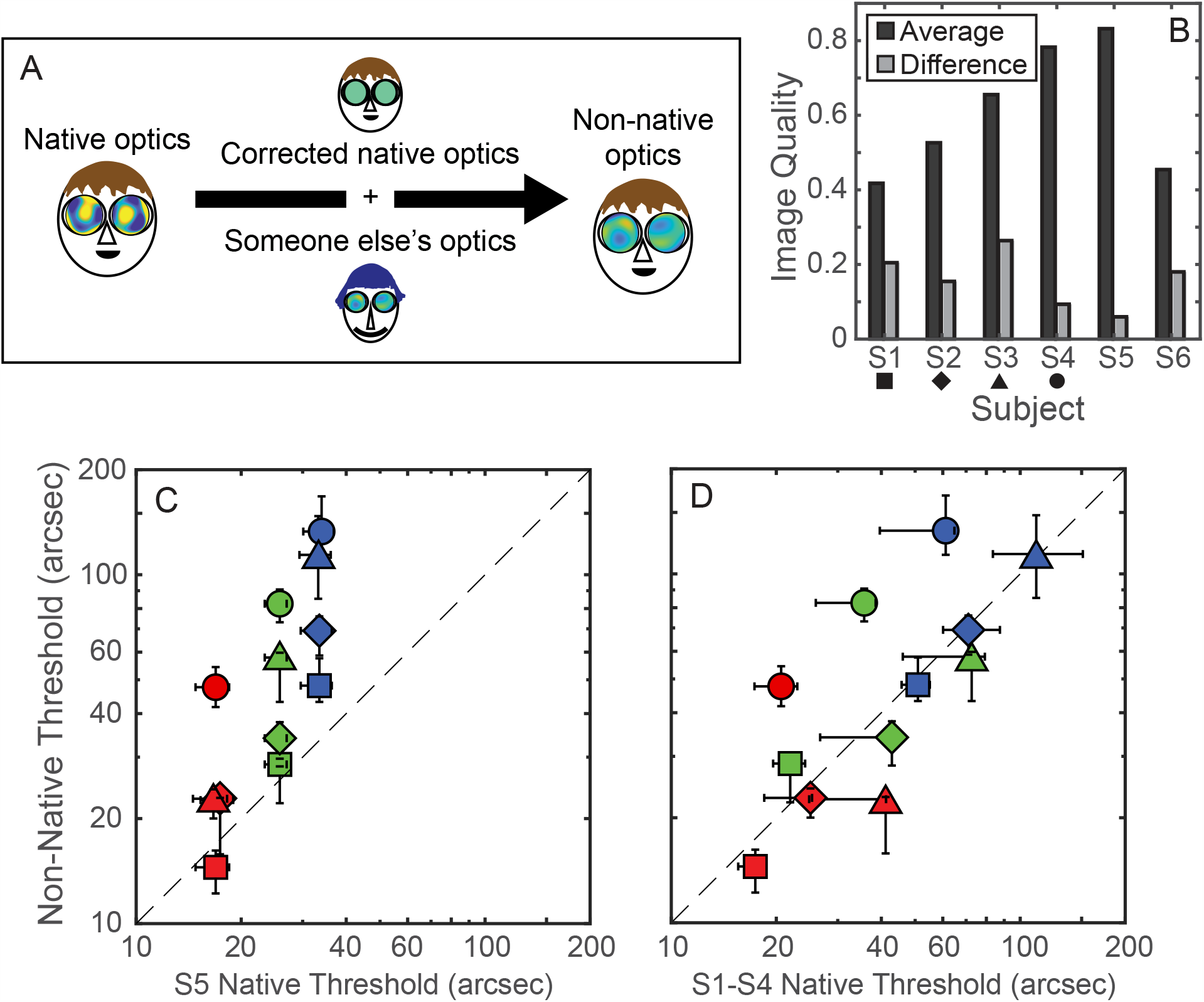
Stereo thresholds with non-native optics. (A) Experimental paradigm. (B) Image quality for each participant. Average quality (black) and inter-ocular difference in quality (gray) are plotted for each of the six participants. We imposed the optics of S5 on the eyes of S1, S2, S3, and S4. We also imposed the optics of S1 on the eyes of S6, and vice versa. (C) Stereo thresholds with non-native optics. The smallest discriminable disparity is plotted for S1, S2, S3, and S4 compared with S5, whose optics were imposed. As in panel (B): ◼ represents S1, ◆ S2, ▲ S3, and ● S4. Colors represent measurements at different corrugation frequencies (red for 1cpd, green for 2cpd, and blue for 3cpd). (D) Comparisons of the stereo thresholds of S1-S4 with their native optics and with S5’s optics. All error bars indicate ±1SD.

We measured the wavefront aberrations of both eyes in six participants (Figure S2). Figure 5B shows average native image quality and inter-ocular difference in quality for each of them. We zeroed in on participant S5 who had the best average quality and the smallest inter-ocular difference in quality. Then, as indicated in Figure 5A, we used the AO system to fully correct the native optics, and simultaneously impose the optics of S5 on participants S1, S2, S3, and S4. By doing so, we made the retinal images the same for all participants, and made the quality of retinal images better than those people were used to experiencing. We then examined stereo acuity when S1-S4 viewed the stimuli with the improved but unfamiliar optics. Based on optical quality alone, we would expect that all four participants viewing the stimuli with the same optics would have the same stereo acuity and that that acuity would be the same as S5’s.

Figure 5C shows the results. The horizontal axis plots stereo thresholds for S5 for corrugation frequencies of 1, 2, and 3cpd. The vertical axis plots thresholds for the other four participants when given the optics of S5. We emphasize that S1-S4 now had the same optics and therefore the same retinal images. The four participants with unfamiliar optics performed more poorly than S5 who was tested with his native optics. Remarkably, S4 had the largest decline even though her image quality (average and interocular difference) was most similar to S5. Although her optical quality was similar to S5’s, her aberration profiles were quite different from his (Figure S2). The fact that she did relatively poorly with S5’s optics indicates that familiarity with one’s optics was important for fine stereopsis.

We also compared stereo thresholds in participants S1-S4’s with their native optics and with the improved but unfamiliar optics of S5. There was no systematic difference (Figure 5D). This finding suggests that there are two offsetting factors that determine how optics affects stereopsis: a benefit from improving optical quality and a detriment from having unfamiliar optics. Notably, the one participant whose stereo acuity did not improve with AO correction (Figure 3B) was among the best performers with the native optics. It was plausible that the optical improvement provided by AO was insufficient to counteract the negative adaptation effects as a result of unfamiliar optics.

To further investigate the importance of familiarity, we swapped the optics between the two participants with the poorest optical quality: S1 and S6. Both participants performed significantly more poorly when given the other person’s optics (particularly at high corrugation frequencies). S1’s thresholds went from 17.4arc sec with her own optics to 19.2arcsec with S6’s optics at 1cpd (a 10.3% increase), from 21.9 to 48.6arcsec at 2cpd (122%), and from 50.7 to 94.7arcsec at 3cpd (87%). S6’s thresholds exhibited even more dramatic changes. His thresholds went from 15.2arcsec with his own optics to 64.9arcsec with S1’s optics at 1cpd (270%), from 25.3 to 158arcsec at 2cpd (525%), and from 56.2arcsec to unmeasurable at 3cpd. These results again illustrate that stereopsis is significantly poorer when viewing stimuli with someone else’s optics even when the optical quality of the participants is equivalent in magnitude. This again strongly suggests that the binocular visual system adapts to particular aspects of retinal images experienced in everyday life.

## Discussion

The adult human visual system has operated for years with the native optics unique to the individual. The optics of our two eyes, in turn, have inscribed their uniqueness on the distributions of light formed on the two retinae: i.e., their metaphorical, unique optical signature. The information embodied in those two images, in turn, is transcribed into neural representations that are utilized in mediating every aspect of visual perception including stereopsis. The present study sheds new light on the consequences of manipulating those optical signatures using AO. Our results disclose that those consequences can be advantageous (i.e., improve stereo acuity) or deleterious (i.e., impair stereo acuity), depending on how closely AO correction conforms to the uniqueness of a given individual’s native optics. The following sections consider these two consequences and their implications for understanding human binocular vision.

### AO-mediated improvement in vision

When aberrations in the habitual optics are corrected in the laboratory using AO, the world temporarily looks noticeably different (e.g., see [35]). Accompanying those changes in visual appearance are significant improvements in visual acuity and contrast sensitivity [5, 23, 36-38]. These improvements are not surprising given blur’s well documented, deleterious impact on resolution [5, 6, 38-43]. Indeed, participants with full AO correction in our experiments exhibited high visual acuity that approached the limit imposed by photoreceptor sampling frequency. Similarly, we found that correcting higher-order aberrations, which are not visually conspicuous in well-corrected eyes, improves stereo acuity, especially with higher frequency modulations in disparity.

The improvement in stereo acuity with AO-corrected optics we observed stands in contrast to results from an earlier study out of our lab [26] suggesting that higher-order aberrations have essentially no impact on stereo acuity. We believe that procedural differences are responsible for the difference in findings. The earlier study used optical phase plates to achieve static correction of higher-order aberrations, whereas the present study used dynamic, real-time AO correction which, unlike the static phase plate, compensates for eye movements and thus mitigates optical effects of pupil/image misalignment that can happen when viewing with a static correction. We are thus confident that the improved AO device and testing procedures employed in this study are responsible for revealing a genuine improvement in stereo acuity attributable to elimination of higher-order aberrations. This, in turn, raises the following question: how does this improvement come about?

### AO improves stereopsis

Our results reveal that levels of stereo acuity achieved with AO exceed those measured when viewing with normal optics. This achievement is remarkable given that the limits of human stereopsis assessed with natural optics already qualifies as a form of hyperacuity: i.e., disparity resolution that exceeds the sampling limits imposed by the photoreceptor mosaic [14, 34, 44-47].

What is the basis of this improvement in stereopsis with AO-correction? It is natural to wonder whether the improvement might arise from more stable, accurate vergence fixation prompted by the enhanced clarity of edge information in the AO-corrected retinal images [48]. We doubt, however, that enhanced vergence stability can account for our results because earlier work on human stereopsis reveals that i) vergence accuracy is unaffected by bandpass spatial-frequency filtering of texture stereograms, a maneuver that mimics blur [49], and ii) fixation disparity (a proxy for vergence error that affects stereopsis [50]) is essentially the same when viewing stereo gratings ranging in spatial frequency from 0.5 to 8cpd [51]. Instead, we are inclined to attribute the improved stereopsis with AO-corrected images to neural processes involved in cortical disparity computation per se.

In this context, then, how does elimination of higher-order optical aberrations enable superior stereo acuity, a cortical process? To tackle that question, we first need to consider the nature of the disparities arising from viewing conditions simulating 3D objects (i.e., a corrugated textured surface in our case) seen from two slightly different viewpoints. There are various ways to conceptualize the nature of those disparities [52, 53]. One is in terms of positional disparities between pairs of matching features. A convenient means for extracting that information would be with location-specific cortical receptive fields that function as spatial-frequency selective neural filters [54]. Another, complementary definition of disparity focuses on disparity in the phase domain [55]. An impetus for this idea comes from physiological studies showing that binocular cortical neurons are sensitive to different phase shifts within pairs of monocular images [56, 57]. Several groups have made the case for the joint involvement of both forms of disparity in mediation of stereopsis [58, 59]. Our aim here is not to critique the different models of stereopsis but, rather, to surmise how AO, through the elimination of higher-order aberrations ordinarily embedded in each eye’s retinal image, might augment the luminance distribution information defining those images.

Higher-order aberrations of the eye’s optics degrade retinal images in three ways: 1) they reduce contrast over a range of spatial frequencies, 2) they eliminate very high spatial frequencies altogether, and 3) they alter phase relationships among spatial frequencies that crucially define spatial information portrayed within images. The disruption of this phase congruency causes a significant loss in key structural elements such as sharp edges that make features hard to match accurately between the eyes. In that way, detecting fine positional disparities become difficult. Correcting the aberrations with AO recovers the phase spectra of low spatial frequencies. Adding the phases of high-frequency components that were unavailable before correction enables phase disparity computation from a larger spectrum of channels. It is plausible that this improvement in both contrast and phase congruency in a broadband stimulus like random-dot stereograms allows the visual system to detect even smaller disparities than those resolved with well-focused normal optics. It is also important to note that further investigation is required to learn whether the binocular system can compensate for the phase disruption through long-term adaptation to the eyes’ native optics and if so, the extent to which the improvement in human stereopsis with perfect optics is compromised by phase adaptation [60, 61].

Putting aside those speculations about the bases of AO’s contribution to improved stereo acuity, we next turn to a second intriguing feature of our results: the consequence of viewing the world through someone else’s optics that was unexpectedly not beneficial despite improvements relative to the participant’s own optics.

### Individual differences in the impact of AO viewing

As noted earlier when discussing the blur paradox, differences in the sharpness of images viewed by the two eyes adversely affects stereo acuity (e.g., [62]) suggesting that matching similar optical quality between the eyes is critical for fine stereopsis. We found that the improvement in stereo acuity measured with AO (i.e., aberration free) was *inversely* related to an individual’s interocular difference in their native, habitually experienced optics. Why would that be the case?

Perhaps a given individual’s visual nervous system adapts to the unique optical profile after a long period of time. This neural adaptation to one’s own optics would improve vision, including stereopsis, under natural, everyday viewing [63], but in a manner specific to the aberrations present in the optics of each eye (e.g. [25]). Viewing with AO correction disrupts that previously stable relation between optical profile and the visual nervous system, with the degree of disruption presumably being greater for those with more pronounced higher-order aberrations. This is just what we found for both visual and stereo acuity (Figs. 2D and 4D). Also consistent with this hypothesis based on neural adaptation were the results from our experiment in which participants were tested while viewing with the optics of another person: this produced poorer performance, even though the non-habitual optics were similar or even better than a person’s own.

This kind of adaptation to blur is not limited to the laboratory. In the eye clinic, it is a common practice not to prescribe full eyewear correction (e.g., for astigmatism) so as to avoid short-term visual discomfort.

### Implications for neural plasticity and clinical relevance

The visual circuitry underlying stereovision was traditionally thought to reach maturity during childhood, beyond which little plasticity remains [64, 65]. However, there have been anecdotal instances of post-pubertal adults recovering stereovision, the most famous of whom is “Stereo-Sue” [12] and more recently Bruce Bridgeman [66]. Using more controlled training paradigms, stereoblind people can also recover stereovision to certain extents [67, 68] implying that the binocular system is more plastic than previously thought. We assume that people adapt because the optics changes gradually throughout the lifespan [69, 70], and yet there appears to be a benefit when viewing with their own native state at the time of testing. We found that thresholds with AO correlate with inter-ocular image quality difference and less so with the average image quality in the native optics, presumably due to long-term adaptation. It is conceivable that the effects are even more substantial in participants with highly aberrated eyes such as those with irregular corneal surface profiles (keratoconus). Keratoconus is a corneal disease that emerges in otherwise normal-sighted individuals during the second or third decade of life and causes very large aberrations that are usually quite different between the two eyes. We observed that these patients even with AO have no or very poor stereopsis. Various advanced vision correction methods [71, 72] that provide supernormal vision are currently available or under development. It is of scientific and clinical interest to address the following question: can normal binocular function be recovered by having the visual system become re-adapted to the new, improved optics over time and if so, how quickly can this neural re-adaptation occur?

## Methods

### Participants

Eight adults participated, including the first and last authors. The gender, age, and eyewear prescription of each individual are provided in Table S1. The participants had eye examinations within the past year and had normal vision while wearing their usual prescription, if any: 20/20 Snellen acuity or better and 40arcsec stereo threshold or better (Randot stereo test). The human participants’ protocol was approved by the University of Rochester Research Review Board. All participants signed an informed consent form before participating. Prior to testing, 1% tropicamide solution was administered to both eyes to produce short-duration mydriasis (pupil dilation) and cycloplegia (paralysis of accommodation).

### Apparatus

The binocular AO system used in this study has been described in detail elsewhere [43]. The apparatus can measure and completely correct and/or manipulate lower- and higher-order optical aberrations while visual performance was being measured with images projected separately to the two eyes. The apparatus consisted of two identical systems, one for each eye. Each had a Shack-Hartmann wavefront sensor that measured the eye’s aberrations from the retinal reflections of a super-luminescent diode at 850nm (Inphenix Inc.). Each wavefront sensor communicated with a deformable mirror (DM-97-15, ALPAO) that controlled the amount and type of optical aberration by conforming its shape to yield the desired wavefront for each eye in real time at 12Hz.

Aberrations were corrected for 6mm pupil diameters while the actual pupil sizes during testing were restricted to 5.8mm using artificial apertures placed at the pupil-conjugate planes. Participants rested their heads on a chin rest and a pair of temple mounts. The rest and mounts could be translated by a 3-axis motorized stage to center both pupils as monitored by a pair of pupil cameras. The same pair of pupil cameras monitored eye movements throughout the experiments to make sure that the visual axis was always aligned to the optical axis of the system. Inter-pupillary distance was set for each participant using a translation stage. Left- and right-eye stimuli were projected on to the retinae by two digital light-processing projectors (DLPDLCR4710EVM-G2, Texas Instrument Inc.), one for each eye. The stimuli were 8.4° wide by 4.7° high spanning 1920×1080 pixels. Each pixel subtended 0.26arcmin. Root mean square wavefront errors as well as image quality during AO correction of individual eyes are provided in Figure S1.

### Visual acuity

Monocular visual acuity was measured with the Tumbling E task [73]. The black letter E was presented for 250ms in one of four orientations on a white background of 120cd/m^2^. Participants indicated the perceived orientation in a four-alternative, forced-choice (4-AFC) response (Figure S3). Auditory feedback was provided for each correct response. Letter size in terms of stroke width ranged from 0.3 to 50arcmin (Snellen 20/6 to 20/1000) and varied over a 40-trial sequence according to the QUEST+ adaptive staircase method [74]. The procedure was repeated three times (120 trials total) to obtain the letter size associated with 72.4% correct using the best-fitting cumulative Weibull function and Bayesian estimation provided by QUEST+.

### Stereo acuity

Binocular stereo thresholds were measured using random-dot stereograms that portrayed a densely textured surface with disparity-defined sinusoidal depth corrugations (Figure 1, 3, S2B; [31]). We used such stimuli because they allow one to eliminate monocular cues and because blur affects the ability to see the depth corrugation [75]. A trial started with the presentation of a fixation target consisting of a small dot and four diagonal lines seen by both eyes along with vertical and horizontal nonius lines seen by one or the other eye. The parts that were seen by both eyes aided accurate alignment of the eyes. The parts seen only by one eye or the other allowed the participant to assess the accuracy of alignment. When the fixation target was properly fused, it looked like one dot and eight lines. Once fusion was achieved, participants initiated stimulus presentation with a key press.

Each dot in the random-dot stereogram was a small bright square (83.5 x 83.5arcsec) on an otherwise dark background. The dot pattern was generated by first populating a hexagonal grid at nodal points spaced 110arcsec apart. Each dot was then randomly displaced from the nodal point with a direction drawn from a uniform distribution ranging from 0 to 2π and a distance from 0 to 55arcsec. Dot density was 180 dots/deg^2^ in a super-Gaussian window (*W*):

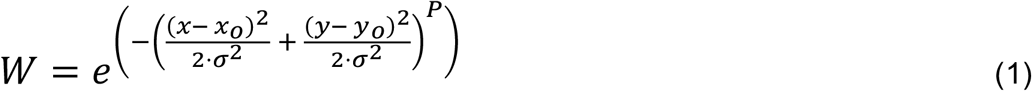

where *P* = 5, and *σ* = 0.5°. The values [*x*_*o*_, *y*_*o*_] are nodal points and [*x, y*] are horizontal and vertical screen coordinates. Edges of the circular window were blended into the background so that the only fusion cues were the random dots themselves.

Left and right images were created from the random-dot pattern by displacing each dot in opposite directions by half its horizontal peak-to-trough disparity (*A*):

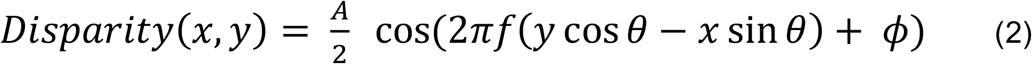

where *f*, *ϕ*, and *θ* are the spatial frequency, phase, and orientation of the disparity-defined sinusoidal corrugation, respectively. Thresholds (the smallest discriminable disparity) were measured for corrugation frequencies of 1, 2, and 3cpd. The corrugation presented on each trial had a random phase between 0 and 2*π* and an orientation of either +10° (slightly anti-clockwise) or −10° (slightly clockwise) from the horizontal. Participants indicated which of the two orientations were presented on each trial, guessing if necessary (2-AFC). Each stimulus was displayed for a maximum of 10s, but participants were instructed to respond as soon as they were confident of their judgment. Most responses were completed under 1s. The peak-to-trough disparity was varied from trial to trial according to the method of constant stimuli. Five disparity amplitudes (determined for each person in pilot testing) were each presented 40 times for a total of 200 trials per condition. We did not present disparity amplitudes that exceeded the disparity-gradient limit [75, 76]. Auditory feedback was provided when a correct response was made. Data for each corrugation frequency were fitted with a cumulative Weibull function using Psignifit [77]. Stereo thresholds were defined as disparities that produced 81.6% correct responses.

### Retinal-image quality

The point-spread function (PSF) represents how a point source of light is blurred by the eye’s aberrations on the retina. PSFs were calculated for the left (*PSF*_*LE*_) and right eyes (*PSF*_*RE*_) of each participant when their lower-order (spherical and cylindrical) refractive errors were corrected (using clinically prescribed eyewear), but their higher-order aberrations were not. The resultant PSFs were a combination of the higher-order aberrations as well as any residual lower-order ones. We quantified retinal-image quality in the following ways. For the visual acuity experiment, we first generated simulated retinal images by convolving an upright 20/20 Snellen E with the eye’s PSF. We then correlated the obtained images with the original perfect images. Specifically, we calculated the two-dimensional cross-correlation and took the maximum as the image-quality value [78]. The metric values can range from −1 to +1 where +1 would mean perfect image quality, unadulterated by aberrations and diffraction, and −1 would indicate anticorrelated image quality. We used the same approach for the stereo experiment, but by convolving a random-dot pattern presented to an eye with the eye’s PSF. We did this for 10 different instances of random-dot patterns. Left-eye image quality (*ImQ*_*LE*_) is:

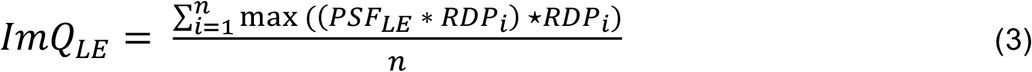

where *RDP* is the random-dot pattern, *n* = 10 represents the 10 instances of *RDP*s, * is 2-D convolution, ⋆ is 2-D cross-correlation, and max(⋆) provides the maximum of the 2-D cross-correlation matrix. Right-eye quality (*ImQ*_*RE*_) was calculated the same way. We also calculated the average image quality (*ImQ*_*Mean*_) defined as the mean of the two eyes’ quality indices across the 10 presentation instances:

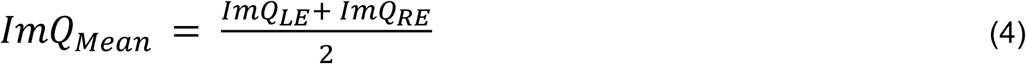

Finally, an index of the interocular difference in image quality (*ImQ*_*IOD*_) was derived by cross-correlating the left and right retinal images, and then subtracting the resultant from unity:

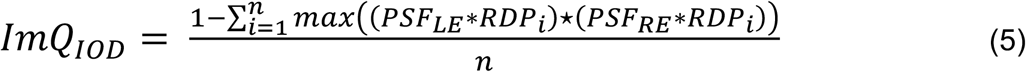

## Acknowledgements

We thank Gregory DeAngelis for helpful comments on the manuscript. This work was supported by NIH Grant EY014999 and Research to Prevent Blindness (RPB).

## Author Contributions

CJN, MSB and GY designed the research. CJN conducted the research. CJN and DT analyzed data. CJN, RB, MSB, DT and GY wrote the paper.

The authors declare no competing interests.

## References

1. Liang, J. and D.R. Williams, Aberrations and retinal image quality of the normal human eye. J Opt Soc Am A Opt Image Sci Vis, 1997. 14(11): p. 2873–83.

2. M., M.J., Molyneux’s Question: Vision, Touch and the Philosophy of Perception. Vol. 1st Edition. 1977, The United States of America: Cambridge University Press.

3. Liang, J., D.R. Williams, and D.T. Miller, Supernormal vision and high-resolution retinal imaging through adaptive optics. Journal of the Optical Society of America A, 1997. 14(11): p. 2884–2892.

4. Marcos, S., et al., Vision science and adaptive optics, the state of the field. Vision Research, 2017. 132: p. 3–33.

5. Yoon, G.Y. and D.R. Williams, Visual performance after correcting the monochromatic and chromatic aberrations of the eye. J Opt Soc Am A Opt Image Sci Vis, 2002. 19(2): p. 266–75.

6. Marcos, S., et al., Influence of adaptive-optics ocular aberration correction on visual acuity at different luminances and contrast polarities. Journal of Vision, 2008. 8(13): p. 1–1.

7. Artal, P., et al., Visual effect of the combined correction of spherical and longitudinal chromatic aberrations. Opt Express, 2010. 18(2): p. 1637–48.

8. Cumming, B.G. and G.C. DeAngelis, The physiology of stereopsis. Annu Rev Neurosci, 2001. 24: p. 203–38.

9. Blake, R. and H.R. Wilson, Neural models of stereoscopic vision. Trends Neurosci, 1991. 14(10): p. 445–52.

10. Barlow, H.B., C. Blakemore, and J.D. Pettigrew, The neural mechanism of binocular depth discrimination. J Physiol, 1967. 193(2): p. 327–42.

11. Wheatstone, C., Contributions to the physiology of visionöPart the first. On some remarkable, and hitherto unobserved, phenomena of binocular vision. Philosophical Transactions of the Royal Society of London, 1838. 128: p. 371–394.

12. Barry, S.R., Fixing My Gaze: A Scientist’s Journey into Seeing in Three Dimensions. 2009, New York, USA: Basic Books.

13. Wood, I.C., Stereopsis with spatially-degraded images. Ophthalmic Physiol Opt, 1983. 3(3): p. 337–40.

14. Westheimer, G. and S.P. McKee, Stereoscopic acuity with defocused and spatially filtered retinal images. Journal of the Optical Society of America, 1980. 70(7): p. 772–778.

15. Odell, N.V., et al., The effect of induced monocular blur on measures of stereoacuity. J aapos, 2009. 13(2): p. 136–41.

16. Nabie, R., et al., Effect of artificial anisometropia in dominant and nondominant eyes on stereoacuity. Can J Ophthalmol, 2017. 52(3): p. 240–242.

17. Schor, C. and T. Heckmann, Interocular differences in contrast and spatial frequency: Effects on stereopsis and fusion. Vision Research, 1989. 29(7): p. 837–847.

18. Fernández, E.J., et al., Impact on stereo-acuity of two presbyopia correction approaches: monovision and small aperture inlay. Biomedical optics express, 2013. 4(6): p. 822–830.

19. Evans, B.J., Monovision: a review. Ophthalmic Physiol Opt, 2007. 27(5): p. 417–39.

20. Schwarz, C., et al., Comparison of binocular through-focus visual acuity with monovision and a small aperture inlay. Biomedical optics express, 2014. 5(10): p. 3355–3366.

21. Porter, J., et al., Monochromatic aberrations of the human eye in a large population. J Opt Soc Am A Opt Image Sci Vis, 2001. 18(8): p. 1793–803.

22. Sawides, L., et al., Dependence of subjective image focus on the magnitude and pattern of high order aberrations. Journal of Vision, 2012. 12(8): p. 4–4.

23. Sabesan, R. and G. Yoon, Neural compensation for long-term asymmetric optical blur to improve visual performance in keratoconic eyes. Invest Ophthalmol Vis Sci, 2010. 51(7): p. 3835–9.

24. Artal, P., et al., Neural compensation for the eye’s optical aberrations. Journal of Vision, 2004. 4(4): p. 4–4.

25. Kompaniez, E., et al., Adaptation to interocular differences in blur. J Vis, 2013. 13(6): p. 19.

26. Vlaskamp, B.N.S., G. Yoon, and M.S. Banks, Human Stereopsis Is Not Limited by the Optics of the Well-Focused Eye. The Journal of Neuroscience, 2011. 31(27): p. 9814–9818.

27. Sabesan, R. and G. Yoon, Visual performance after correcting higher order aberrations in keratoconic eyes. Journal of Vision, 2009. 9(5): p. 6–6.

28. Ahnelt, P.K., The photoreceptor mosaic. Eye (Lond), 1998. 12 (Pt 3b): p. 531–40.

29. Jonas, J.B., U. Schneider, and G.O. Naumann, Count and density of human retinal photoreceptors. Graefes Arch Clin Exp Ophthalmol, 1992. 230(6): p. 505–10.

30. Kane, D., P. Guan, and M.S. Banks, The limits of human stereopsis in space and time. J Neurosci, 2014. 34(4): p. 1397–408.

31. Tyler, C.W., Depth perception in disparity gratings. Nature, 1974. 251(5471): p. 140–142.

32. Bosten, J.M., et al., A population study of binocular function. Vision Res, 2015. 110(Pt A): p. 34–50.

33. Peterzell, D.H., et al., Thresholds for sine-wave corrugations defined by binocular disparity in random dot stereograms: Factor analysis of individual differences reveals two stereoscopic mechanisms tuned for spatial frequency. Vision Research, 2017. 141: p. 127–135.

34. Westheimer, G. and S.P. McKee, Stereogram design for testing local stereopsis. Invest Ophthalmol Vis Sci, 1980. 19(7): p. 802–9.

35. Artal, P., et al., Neural compensation for the eye’s optical aberrations. J Vis, 2004. 4(4): p. 281–7.

36. Sabesan, R., A. Barbot, and G. Yoon, Enhanced neural function in highly aberrated eyes following perceptual learning with adaptive optics. Vision Research, 2017. 132: p. 78–84.

37. Rossi, E.A., et al., Visual performance in emmetropia and low myopia after correction of high-order aberrations. Journal of Vision, 2007. 7(8): p. 14–14.

38. Schwarz, C., et al., Binocular visual acuity for the correction of spherical aberration in polychromatic and monochromatic light. Journal of Vision, 2014. 14(2): p. 8–8.

39. Campbell, F.W. and D.G. Green, Optical and retinal factors affecting visual resolution. The Journal of physiology, 1965. 181(3): p. 576–593.

40. Liang, J., D.R. Williams, and D.T. Miller, Supernormal vision and high-resolution retinal imaging through adaptive optics. J Opt Soc Am A Opt Image Sci Vis, 1997. 14(11): p. 2884–92.

41. Li, S., et al., Effects of Monochromatic Aberration on Visual Acuity Using Adaptive Optics. Optometry and Vision Science, 2009. 86(7).

42. Rocha, K.M., et al., Enhanced visual acuity and image perception following correction of highly aberrated eyes using an adaptive optics visual simulator. J Refract Surg, 2010. 26(1): p. 52–6.

43. Sabesan, R., L. Zheleznyak, and G. Yoon, Binocular visual performance and summation after correcting higher order aberrations. Biomed Opt Express, 2012. 3(12): p. 3176–89.

44. McKee, S.P., The spatial requirements for fine stereoacuity. Vision Res, 1983. 23(2): p. 191–8.

45. Stevenson, S.B., L.K. Cormack, and C.M. Schor, Hyperacuity, superresolution and gap resolution in human stereopsis. Vision Research, 1989. 29(11): p. 1597–1605.

46. Westheimer, G., Editorial: Visual acuity and hyperacuity. Invest Ophthalmol, 1975. 14(8): p. 570–2.

47. Westheimer, G., Cooperative neural processes involved in stereoscopic acuity. Exp Brain Res, 1979. 36(3): p. 585–97.

48. Otero-Millan, J., S.L. Macknik, and S. Martinez-Conde, Fixational eye movements and binocular vision. Frontiers in integrative neuroscience, 2014. 8: p. 52–52.

49. Mowforth, P., J.E. Mayhew, and J.P. Frisby, Vergence eye movements made in response to spatial-frequency-filtered random-dot stereograms. Perception, 1981. 10(3): p. 299–304.

50. Ukwade, M.T., H.E. Bedell, and R.S. Harwerth, Stereopsis is perturbed by vergence error. Vision Res, 2003. 43(2): p. 181–93.

51. Harwerth, R.S., E.L. Smith, 3rd, and J. Siderov, Behavioral studies of local stereopsis and disparity vergence in monkeys. Vision Res, 1995. 35(12): p. 1755–70.

52. Qian, N. and S. Mikaelian, Relationship Between Phase and Energy Methods for Disparity Computation. Neural Computation, 2000. 12(2): p. 279–292.

53. Blake, R. and H. Wilson, Binocular vision. Vision Res, 2011. 51(7): p. 754–70.

54. Fleet, D.J., H. Wagner, and D.J. Heeger, Neural encoding of binocular disparity: Energy models, position shifts and phase shifts. Vision Research, 1996. 36(12): p. 1839–1857.

55. Lappin, J.S., What is binocular disparity? Frontiers in psychology, 2014. 5: p. 870–870.

56. Ohzawa, I., G.C. DeAngelis, and R.D. Freeman, Stereoscopic depth discrimination in the visual cortex: neurons ideally suited as disparity detectors. Science, 1990. 249(4972): p. 1037–41.

57. Tsao, D.Y., B.R. Conway, and M.S. Livingstone, Receptive Fields of Disparity-Tuned Simple Cells in Macaque V1. Neuron, 2003. 38(1): p. 103–114.

58. Lappin, J.S. and W.D. Craft, Definition and detection of binocular disparity. Vision Res, 1997. 37(21): p. 2953–74.

59. Read, J.C.A. and B.G. Cumming, Sensors for impossible stimuli may solve the stereo correspondence problem. Nature Neuroscience, 2007. 10(10): p. 1322–1328.

60. Barbot, A., et al., Neural adaptation to optical aberrations through phase compensation. Journal of Vision, 2020. 20(11): p. 1130–1130.

61. Barbot, A., et al., Long-term adaptation to ocular aberrations alters visual processing of spatial frequency information. Journal of Vision, 2016. 16(12): p. 554–554.

62. Lam, A.K.C., et al., Effect of naturally occurring visual acuity differences between two eyes in stereoacuity. Ophthalmic and Physiological Optics, 1996. 16(3): p. 189–195.

63. Webster, M.A., M.A. Georgeson, and S.M. Webster, Neural adjustments to image blur. Nat Neurosci, 2002. 5(9): p. 839–40.

64. Daw, N.W., Critical Periods and Amblyopia. Archives of Ophthalmology, 1998. 116(4): p. 502–505.

65. Banks, M.S., R.N. Aslin, and R.D. Letson, Sensitive period for the development of human binocular vision. Science, 1975. 190(4215): p. 675.

66. Bridgeman, B., Restoring Adult Stereopsis: A Vision Researcher’s Personal Experience. Optometry and Vision Science, 2014. 91(6).

67. Ding, J. and D.M. Levi, Recovery of stereopsis through perceptual learning in human adults with abnormal binocular vision. Proceedings of the National Academy of Sciences, 2011. 108(37): p. E733–E741.

68. Vedamurthy, I., et al., A dichoptic custom-made action video game as a treatment for adult amblyopia. Vision Res, 2015. 114: p. 173–87.

69. Athaide, H.V., M. Campos, and C. Costa, Study of ocular aberrations with age. Arq Bras Oftalmol, 2009. 72(5): p. 617–21.

70. Amano, S., et al., Age-related changes in corneal and ocular higher-order wavefront aberrations. Am J Ophthalmol, 2004. 137(6): p. 988–92.

71. Marsack, J.D., et al., Wavefront-guided scleral lens correction in keratoconus. Optom Vis Sci, 2014. 91(10): p. 1221–30.

72. Sabesan, R., et al., Wavefront-guided scleral lens prosthetic device for keratoconus. Optom Vis Sci, 2013. 90(4): p. 314–23.

73. Lovie-Kitchin, J.E., Validity and reliability of visual acuity measurements. Ophthalmic Physiol Opt, 1988. 8(4): p. 363–70.

74. Watson, A.B., QUEST+: A general multidimensional Bayesian adaptive psychometric method. Journal of Vision, 2017. 17(3): p. 10–10.

75. Banks, M.S., S. Gepshtein, and M.S. Landy, Why Is Spatial Stereoresolution So Low? The Journal of Neuroscience, 2004. 24(9): p. 2077.

76. Burt, P. and B. Julesz, A disparity gradient limit for binocular fusion. Science, 1980. 208(4444): p. 615–7.

77. Wichmann, F.A. and N.J. Hill, The psychometric function: I. Fitting, sampling, and goodness of fit. Perception & Psychophysics, 2001. 63(8): p. 1293–1313.

78. Zheleznyak, L., et al., Impact of corneal aberrations on through-focus image quality of presbyopia-correcting intraocular lenses using an adaptive optics bench system. J Cataract Refract Surg, 2012. 38(10): p. 1724–33.

