## Supplemental figures for "Seeing the World Like Never Before: Human stereovision through perfect optics"

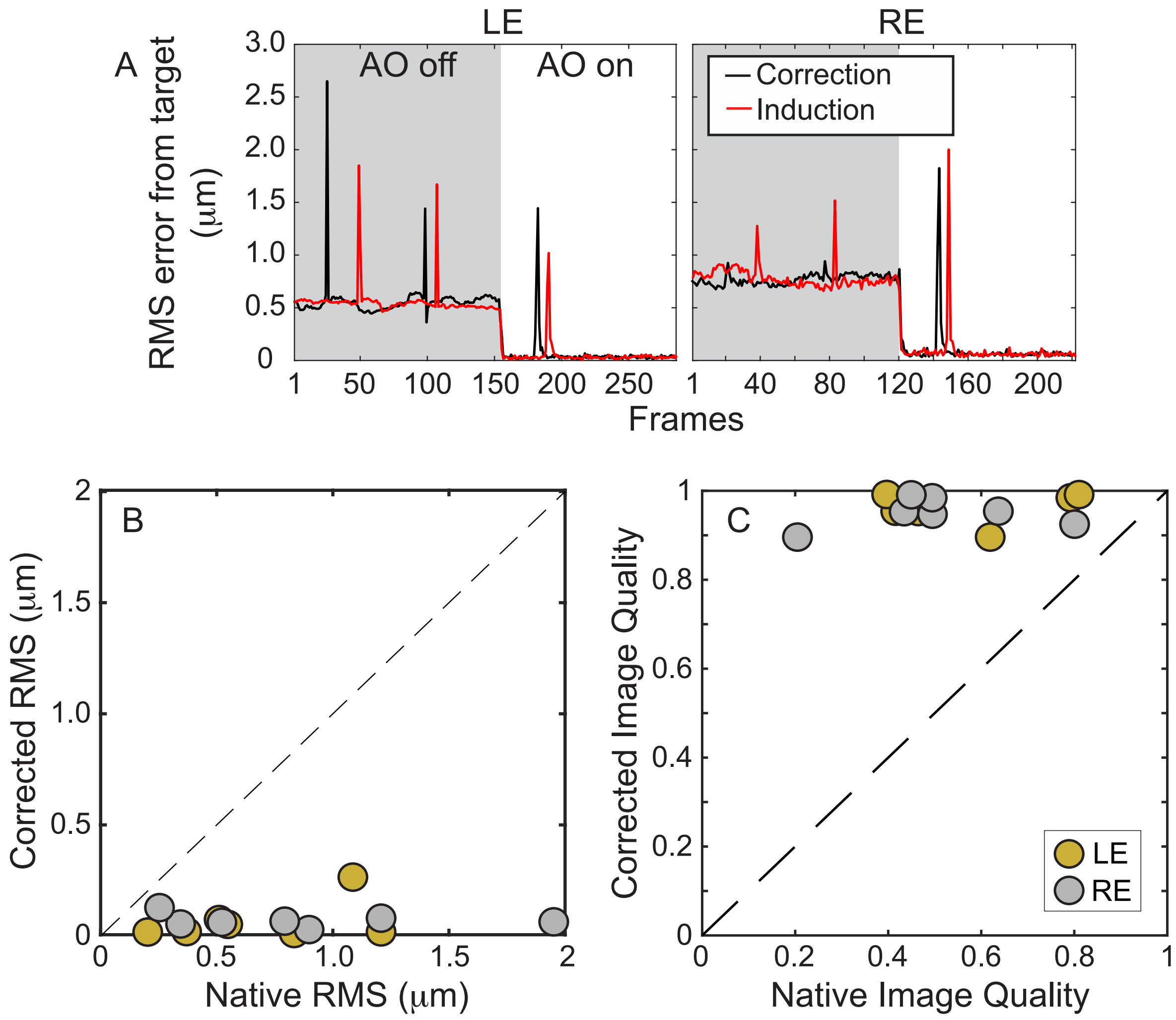


**Fig. S1. Native vs AO optical quality at 5.8mm pupil size.** (A) Time course changes to the optics of both eyes in an example participant (S1) as a result of eye movements. Grey zones indicate the native optical quality without the AO switched on. White zones are when the AO was switched on to have the participant either fully corrected (black traces) or additionally induced with non-native optics (red trace). RMS error are deviations from the target wavefronts; the target being a perfectly flat wavefront in the case of full AO correction or having S5's wavefront during induction with the non-native optics. The RMS error of each eye was calculated by taking the root of the squared and summed Zernike coefficients up to the fifth order measured with the wavefront sensing arm of the AO. Tip and tilt were set to zero. Spikes were instances of blinking during which AO correction paused. Since the AO was run at 12Hz, each frame that comprised of a single wavefront measurement and deformable mirror correction lasted for 5s. (B) & (C) Optical quality with and without AO correction across all participants as given by the RMS error of the wavefront (B) or image quality from the cross-correlation metric as given in Equation 3 of the Methods (C).These plots illustrate the large inter-subject variability in the native optical quality that became comparable during AO correction.

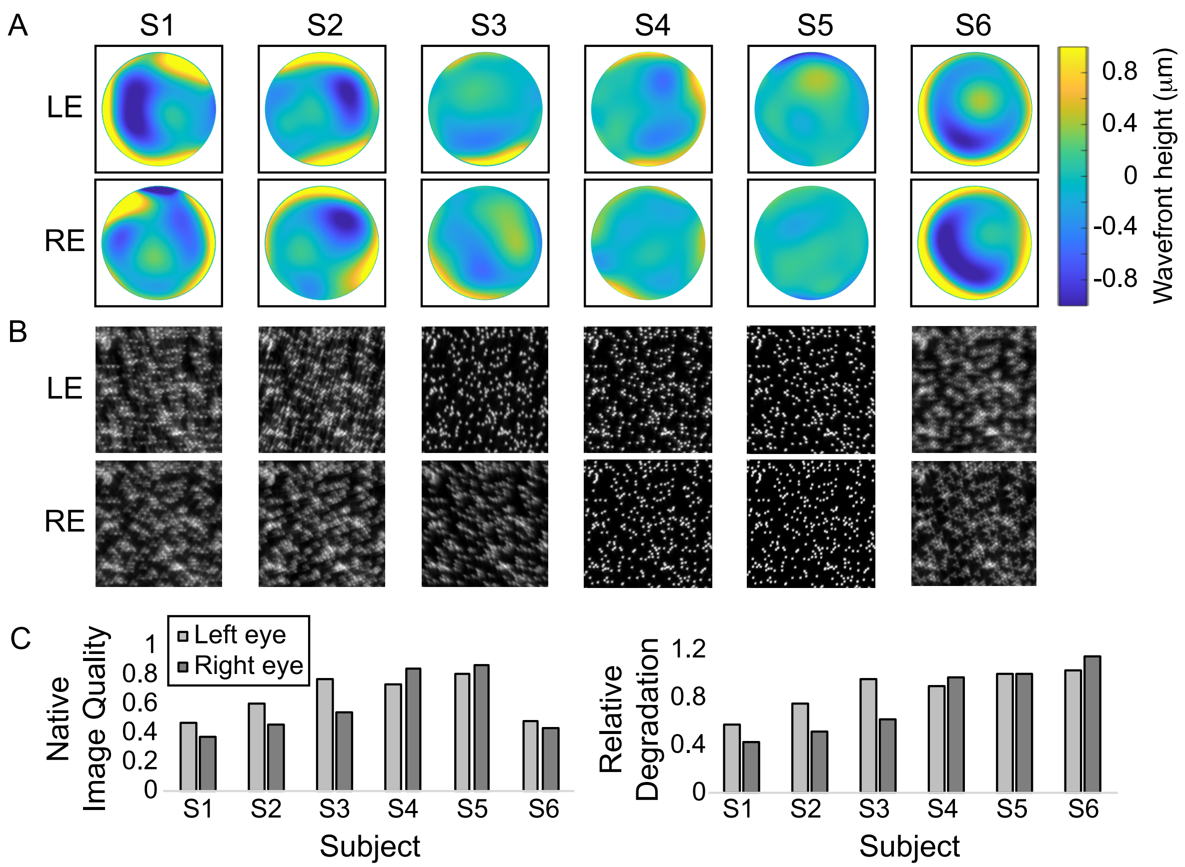


**Figure S2. Native optical profiles of the participants whose stereo-thresholds were measured with S1's aberrations.** (A) Wavefront profiles of the left (top row) and right (bottom row) eyes. Color deviation indicate increasing severity of the aberrations. Wavefront profiles of individual eyes were reconstructed by summing the averages of 100 frames of the Zernike coefficients without blinks up to the fifth order and with tip and tilt set to zero (see Fig. S1A). (B) Simulated retinal images of individual eyes when viewing a random dot pattern. The simulated random dot images were obtained by convolving the example random dot image by the respective eyes' PSF (see Methods). (C) Image quality of individual eyes calculated from Equation 3. Briefly, the simulated retinal image was cross-correlated with the original example image. The retinal image quality was taken as the maximum of the two-dimensional cross-correlation coefficient. (D) Relative image quality degradation from the non-habitual aberrations that was imposed, which in the case of S1-S5, was calculated by dividing the individual eye's image quality by S5's. S6 was divided by S1's image quality. Values below 1 indicate habitual retinal image quality that was worse than the imposed.

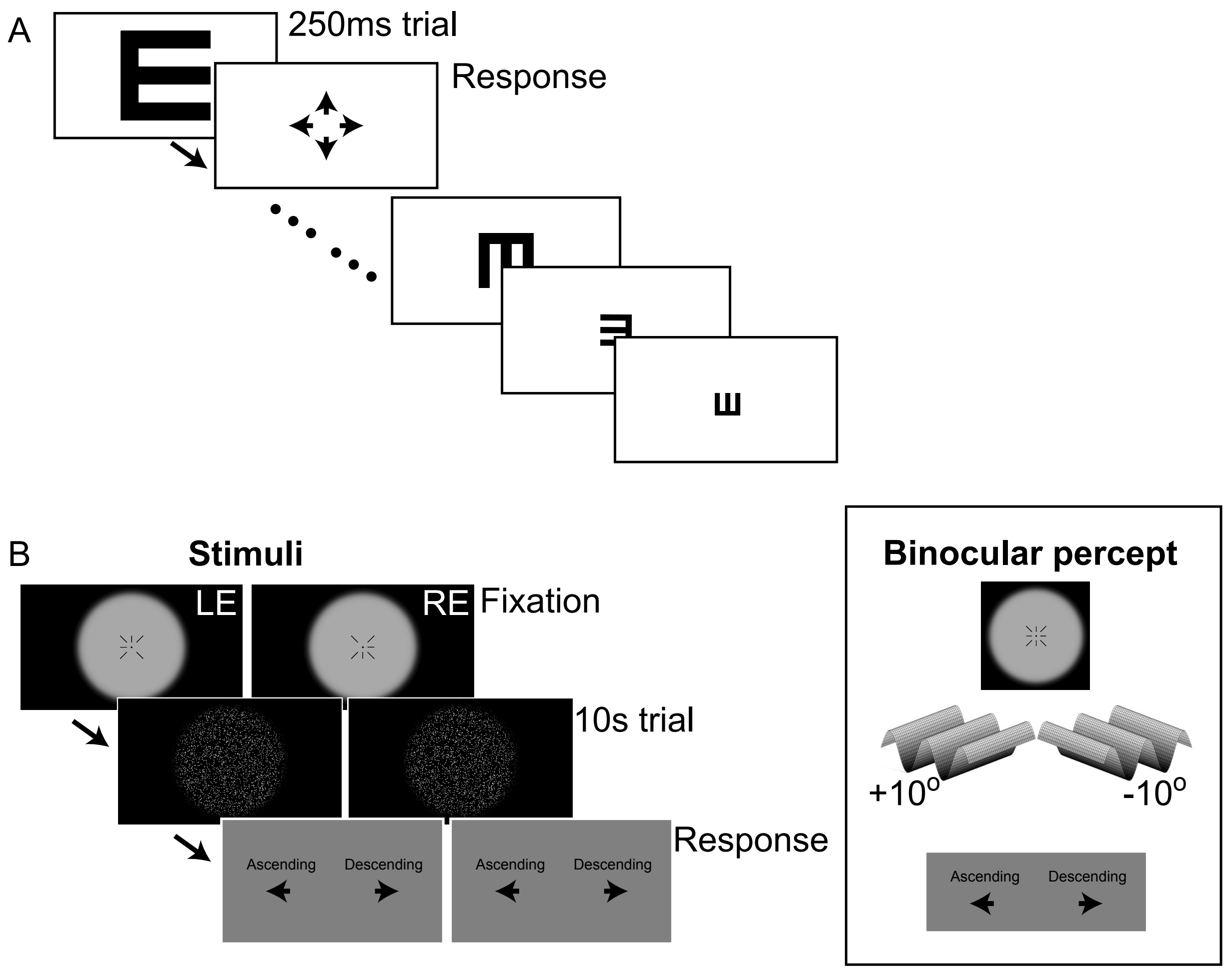


**Fig. S3. Experimental stimuli and procedure (see Methods for details). (**A) Monocular visual acuity measured with the Tumbling E method. The letter E was presented in one of the four depicted orientations on every trial. Participants covered the untested eye and viewed the stimulus monocularly. After 250ms, the letter E stimulus disappeared, initiating a response of the orientation of letter "E" that had just been presented. Letter size was progressively made smaller using a QUEST+ adaptive staircase procedure until the orientation was no longer discernible. Each block provided a threshold measurement that was repeated three times per eye of each participant. Visual acuity of individual eyes was the average of the three. (B) Sine wave corrugation stereoacuity. Stimuli were presented simultaneously to both eyes as depicted in the left panel. The right panel show the binocular appearance of the respective presentations. Every trial was preceded by a fixation target of four diagonal lines and a central dot presented dioptically, and a pair of cardinal nonius lines presented dicoptically. Participants viewed the fixation screens until proper fusion was achieved, as signaled by the appearance of eight lines surrounding the central dot (right panel). Then, the trial was initiated with a keypress. Stimuli were generated according to Equations 1 and 2 of the Methods section. When fused, the stimulus in each trial was a sinusoidal wave pattern modulated in depth either oriented +10° or -10° from the horizontal. Participants were given 10s to view the stimulus. A response screen replaced the stimulus upon termination of the 10s. Subjects were free to respond anytime during or after the stimulus presentation, but instructed to do so as soon as they perceived the corrugation to be ascending (+10°) or descending (-10°).

**Table S1. Participant details**. A total of eight normally sighted adults participated in this study. The mean age was 29.4±11.2 years. All participants were emmetropic to moderately myopic (<-5D), and either had no or mild astigmatism (<1D). All routinely wore well corrected prescription eyewear in everyday viewing if necessary, as given in the last two columns in diopters. Hence, the residual aberrations that we measured were mainly a result of higher-order aberrations. Both eyes of all participants were screened negative for ocular disease, intraocular scatter and binocular vision anomalies. Informed consent was obtained prior to experimental measurements.

| Participant | Age (years) | Gender | Condition | Left eye Prescription (SPH/CYL/axis) | Right eye Prescription (SPH/CYL/axis) |
|---|---|---|---|---|---|
| S1 (author) | 39 | F | Emmetrope | Plano | Plano |
| S2 | 27 | F | Myope | -3.25/0/0 | -3.25/0/0 |
| S3 | 22 | F | Myope | -1.00/-0.5/16 | -1.25/0/0 |
| S4 (only Figure6) | 24 | F | Myope | -3.75/-0.75/40 | -4.75/0/0 |
| S5 | 21 | M | Emmetrope | Plano | Plano |
| S6 (author) | 53 | M | Emmetrope | 0/-0.75/0 | -0.25/0.75/0 |
| S7 | 28 | M | Emmetrope | Plano | Plano |
| S8 | 21 | F | Myope | -2.25/0/0 | -2.25/-0.75/175 |
